# High resolution live cell imaging to define ultrastructural and dynamic features of the halotolerant yeast *Debaryomyces hansenii*

**DOI:** 10.1101/2024.03.01.582977

**Authors:** Martha S.C. Xelhuantzi, Daniel Ghete, Amy Milburn, Savvas Ioannou, Phoebe Mudd, Grant Calder, José Ramos, Peter J. O’Toole, Paul G. Genever, Chris MacDonald

## Abstract

Although some budding yeasts have proved tractable and intensely studied models, others are more recalcitrant. *Debaryomyces hansenii*, an important yeast species in food and biotechnological industries with curious physiological characteristics, has proved difficult to manipulate genetically and remains poorly defined. To remedy this, we have combined live cell fluorescent dyes with high resolution imaging techniques to define the sub-cellular features of *D. hansenii*, such as the mitochondria, nuclei, vacuoles and the cell wall. Using these tools, we define biological processes like the cell cycle, organelle inheritance and different membrane trafficking pathways of *D. hansenii* for the first time. Beyond this, reagents designed to study *Saccharomyces cerevisiae* proteins were used to access proteomic information about *D. hansenii*. Finally, we optimised the use of label free holotomography to image yeast, defining the physical parameters and visualising sub-cellular features like membranes and vacuoles. Not only does this work shed light on *D. hansenii* but this combinatorial approach serves as a template for how other cell biological systems, which are not amenable to standard genetic procedures, can be studied.

## INTRODUCTION

*Debaryomyces hansenii* is a halophilic and osmotolerant yeast species that has been of interest for basic and applied research due to its unique ability to withstand high osmotic pressure, high salinity, and low water activity (Breuer and Harms, 2006). This yeast is notably recognized for its involvement in the ripening processes of meats and cheeses, as well as its role in the synthesis of valuable chemicals like xylitol and riboflavin (Prista et al., 2016; Yaguchi et al., 2017). Despite its potential, some structural and physiological peculiarities together with the slow development of genetic tools for *D. hansenii* have limited its scientific use and biotechnological potential. Early studies considered *D. hansenii* as a possible emergent pathogen (St-Germain and Laverdière, 1986; Wagner et al., 2005), but this was most probably due to misidentification of the strains, since this yeast is phylogenetically related to some *Candida* species and distinguishing between species was previously difficult (Desnos-Ollivier et al., 2008; Nishikawa et al., 1996; Nishikawa et al., 1997).

Beyond this, studies in *D. hansenii* have been further hindered due to its intractability to typical techniques used for other yeast species (e.g. disruption of the cell wall), potentially low fermentive capacity, and the ability to adapt and grow in the presence of different antibiotics (Ruiz-Pérez et al., 2023). Moreover, and importantly, genetic heterogeneity and distinct chromosome polymorphism has been argued to explain contradictory findings (Huang et al., 2021; Petersen and Jespersen, 2004). The slow development of genetic tools for *D. hansenii* has been affected by several factors. One of the biggest drawbacks has been the absence of efficient transformation protocols until recent years (Minhas and Biswas, 2019; Minhas et al., 2009). Furthermore, the availability of only a few circular plasmids capable of propagation in this yeast posed a major challenge, as these plasmids relied on auxotrophic markers and were applicable only to specific laboratory strains (Maggi and Govind, 2004; Ricaurte and Govind, 1999). To complicate matters, the genetic engineering of *D. hansenii* has been severely limited due to its membership in the CTG clade, which encodes serine instead of leucine and the preference for non-homologous end joining in integration of DNA in chromosomes, making it extremely difficult to introduce precise alterations to the genome through homology-directed repair (Kawaguchi et al., 1989; Prista et al., 2016). Nevertheless, recent advances have been made in the genetic engineering of *D. hansenii*, particularly with the development of a CRISPR/Cas9 method that facilitates efficient oligo-mediated gene editing (Spasskaya et al., 2021; Strucko et al., 2021). Although that method is not yet widely used, it is plausible that this progress will be significant as it allows for precise gene editing, which is essential for the development of *D. hansenii* as a cell factory for various biotechnological applications (Navarrete et al., 2022).

Due to the potential of *D. hansenii* it is unfortunate that the above-described barriers have limited understanding the ultrastructural organisation and physiological regulation of this yeast species. Beyond some original electron microscopy analyses (Gezelius and Norkrans, 1970; Rij and Veenhuis, 1975), little has been learnt about organelle organisation since; even less is known about dynamic processes in living *D. hansenii* cells. This is particularly frustrating given that other yeast species, such as the budding yeast *Saccharomyces cerevisiae*, have an extensive arsenal of tools and reagents that have been used to define eukaryotic cell biology and drive industrial applications (Duina et al., 2014; Mattanovich et al., 2014). To overcome this paucity, we have optimised the use of fluorescent dyes with super resolution microscopy, in addition to label free holotomgraphy, to elucidate various cell biological aspects of *D. hansenii*. Our work reveals many physiological details, including a 3D segmentation-based identification of stages cell cycle stage, intracellular organisation of different organelles, trafficking through the vacuolar degradation and endosomal recycling pathways, and observations of organelle inheritance.

## RESULTS

### *Comparison of Debaryomyces hansenii* and *Saccharomyces cerevisiae*

Initial brightfield imaging experiments using Differential Interference Contrast (DIC) confirmed that *Debaryomyces hansenii* are markedly smaller in size than *Saccharomyces cerevisiae* (**Figure 1A**). To quantify this difference, measurements of cellular area segmented from brightfield micrographs showed a mean area of 17.2 ± 5.8 μm^2^ for *S. cerevisiae* compared with 10.2 ± 5.8 μm^2^ for *D. hansenii* (**Figure 1B**). We have previously noted that this type of analysis from dividing cells results in a wide range of areas (Laidlaw et al., 2022a), so we sampled more than 600 cells per condition. To underscore this difference, we cultured a strain of *S. cerevisiae* expressing a GFP tagged flavodoxin-like protein Ycp4 (Grandori and Carey, 1994), which localises to eisosomes to promote quiescence (Megarioti et al., 2023). GFP-Ycp4 expressing *S. cerevisiae* cells were cultured in parallel with unlabelled *D. hansenii* then mixed. Confocal imaging of the co-culture, with *S. cerevisiae* cells identified by expression of GFP-Ycp4, was used to compare to the relatively smaller *D. hansenii* cells in the same micrographs (**Figure 1C**).

**Figure 1:**
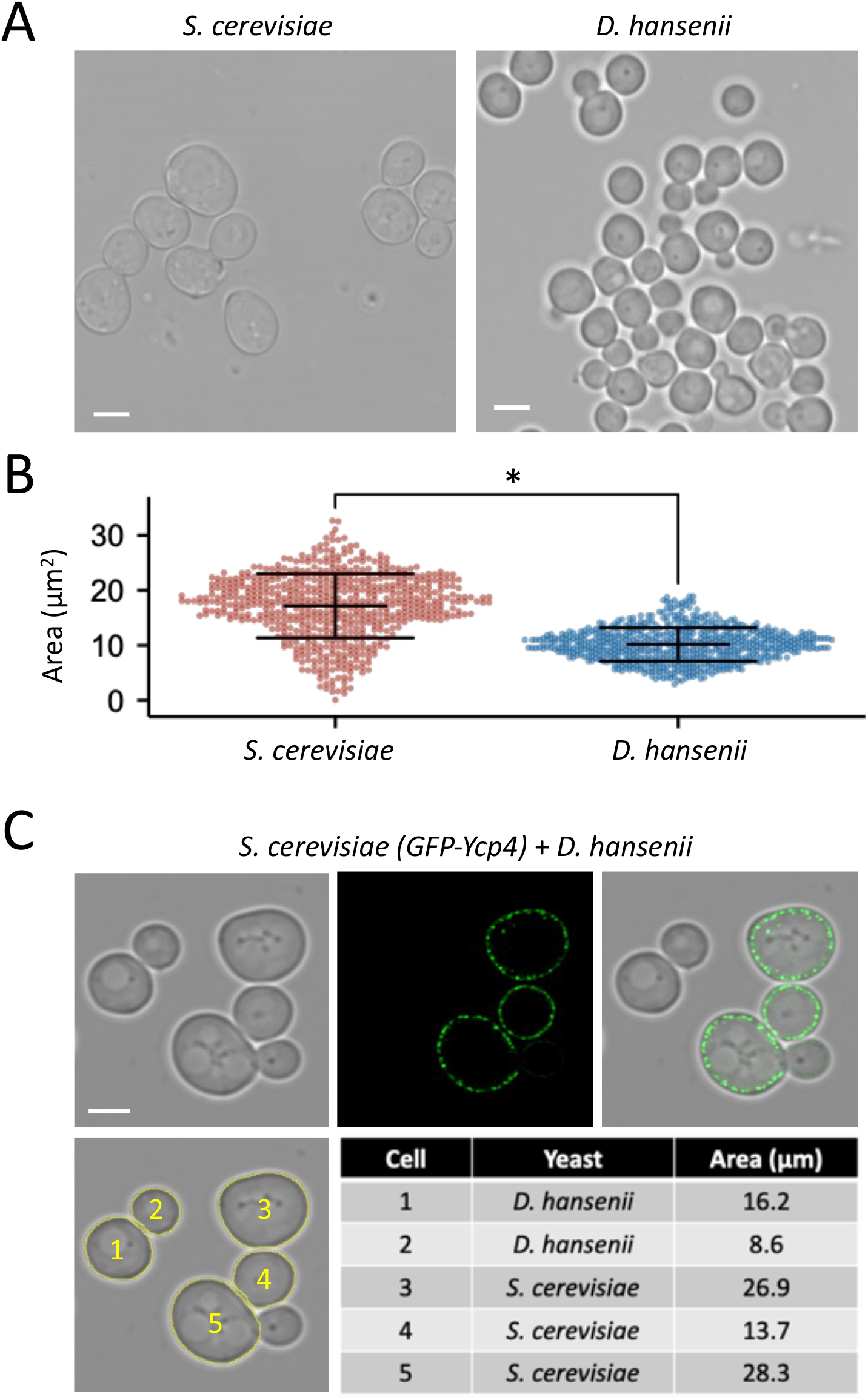
Size comparison between *D. hansenii* and *S. cerevisiae*. **A)** Indicated yeast species were grown to mid-log phase before preparation for brightfield imaging. **B)** Cells from (**A**) were segmented using the Cell Magic Wand Plugin and area measurements calculated (n > 600 cells for each condition) with individual data points displayed and standard deviation shown in error bars. **C)** *S. cerevisiae* cells expressing GFP tagged Ycp4 were cultured to log phase alongside unlabelled *D. hansenii* cells before cultures were mixed and prepared for confocal microscopy. Individual cell segmentation is shown in yellow (lower) and representative area measurements shown in the table. Statistical significance was determined using Student’s *t*-test, with * indicating p < 0.05. Scale bar = 3μm

In an effort to analyse cellular aspects of yeast without using fluorescent proteins we first optimised holotomography for imaging of *S. cerevisiae* cells based on their relative refractive index (**Figure 2A**). The yeast cells could be clearly visualised from this rapid imaging technique, and using the 3D analysis software individual cells could be identified and segmented for downstream processing (**Figure 2B**). We imaged large fields of view of cultured *S. cerevisiae* and *D. hansenii* cells and segmented to create holotomograms for individual cells. We manually categorised these cells based on cell cycle stage, with cells assigned to groups of single cells with no buds, cells with small daughter cells budding, and cells later in the cell division process with large buds (**Figure 2C**). Encouragingly, the described holotomography analysis could be performed in rich YPD media, which is essentially incompatible with fluorescence microscopy due to its auto-fluorescence (Brouwer et al., 2020). Holotomography allows for a range of physical parameters to be measured effortlessly from these segmented cells. For example, estimates of dry mass, volume and length confirmed *S. cerevisiae* cells are consistently larger than the *D. hansenii* counterparts at the same stage of the cell cycle (**Figure 2D**). To ascertain stages of the cell cycle in an automated manner, we compared sphericity measurements from the pre-defined categories of *S. cerevisiae* single cells (e.g. a mother cell in G1 phase or a newly budded daughter cell) or cells in the active process of budding (**Figure 2E**). Single cells have sphericity >0.84 and dividing cells have sphericity <0.82. As with other parameters, when the sphericity of non-budding *S. cerevisiae* cells were compared between YPD and SC media, no significant difference was observed, supporting the notion that holotomography can be performed equally well in rich YPD culture media. We then confirmed cell cycle stage of *D. hansenii* using sphericity with single cells >0.75 and diving cells <0.72 (**Figure 2F**).

**Figure 2:**
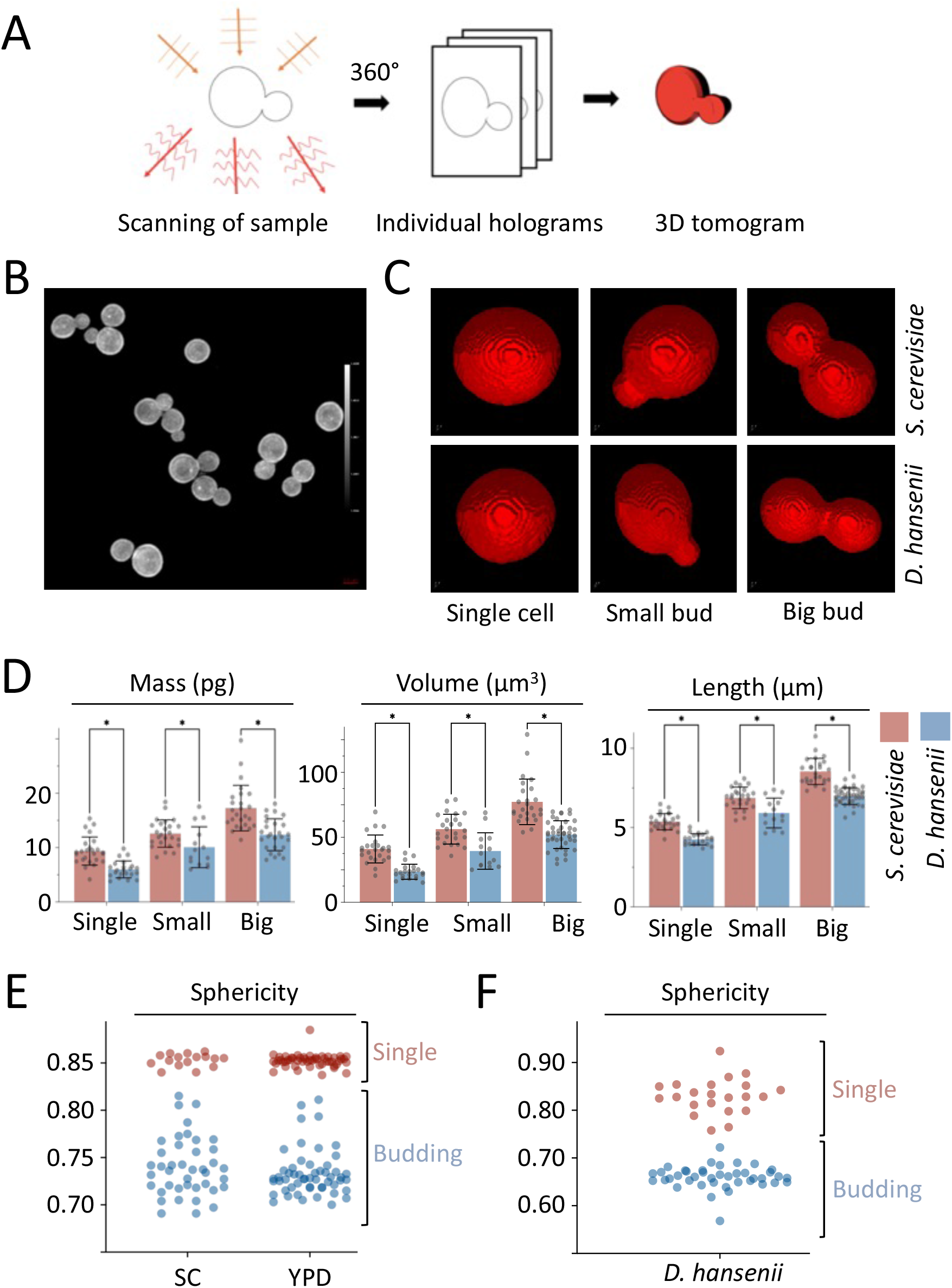
Holotomography for label free live cell imaging of budding yeast. **A)** Schematic cartoon showing holotomography process of rapidly acquired tomograms used to generate 3D tomograms.**B)** Representative field of view showing refractive index based imaging of *S. cerevisiae* cells grown to mid-log phase.**C)** Example 3D tomograms generated from holotomographic imaging of indicated species from single cells or cells early (small bud) or late (big bud) in the cell division process. **D)** Indicated cells from *S. cerevisiae* (red) and *D. hansenii* (blue) were segmented manually and analysed using TomoSuite analysis tools, providing values for mass (pg), volume (μm^3^), and length (μm). n= >71 cells for each yeast species). Statistical significance was determined using Student’s *t*-test, with * indicating p < 0.05. **E - F)** The Tomocube analysis software was used to calculate the sphericity of *S. cerevisiae* in both YPD and SC media **(E)** and *D. hansenii* **(F)**.

Although *D. hansenii* clearly exhibits different physical and physiological features to *S. cerevisiae*, we considered analysing *D. hansenii* proteins using antibodies raised against *S. cerevisiae* proteins. For this, we used antibodies that had previously been used to recognise: the E3 ubiquitin ligase Rsp5 (Stamenova et al., 2004), the eisosme factor Pil1 (Walther et al., 2007), and the endosomal sorting complex required for transport (ESCRT) associated protein Ist1 (Tan et al., 2015). The respective orthologues were identified using protein-protein BLASTP (Camacho et al., 2009) against *Debaromyces hansenii* (taxid:4959): Rsp5 / DEHA2D03718; Pil1 / DEHA2F01694; Ist1 / DEHA2C12408 (**Figure 3A**). Most *D. hansenii* counterparts are similar in predicted molecular weight but with a range of sequence similarity. However, even Ist1 with lowest homology shares many regions across diverse yeast species (**Figure 3B - 3C**). Normalising lysates based on optical density did not create equal loading of lysates, presumably due to difference in cellular size, but loading 3-fold more (3x) *D. hansenii* lysate provided relatively similar proteome profiles, albeit with very distinct protein enrichment profiles (**Figure 3D**). Using this harvesting process, we confirmed that orthologues of Rsp5, Pil1 and Ist1 could all be detected at predicted molecular weights, although levels of the *D. hansenii* Rsp5 orthologue DEHA2D03718 was comparatively low (**Figure 3E**).

**Figure 3:**
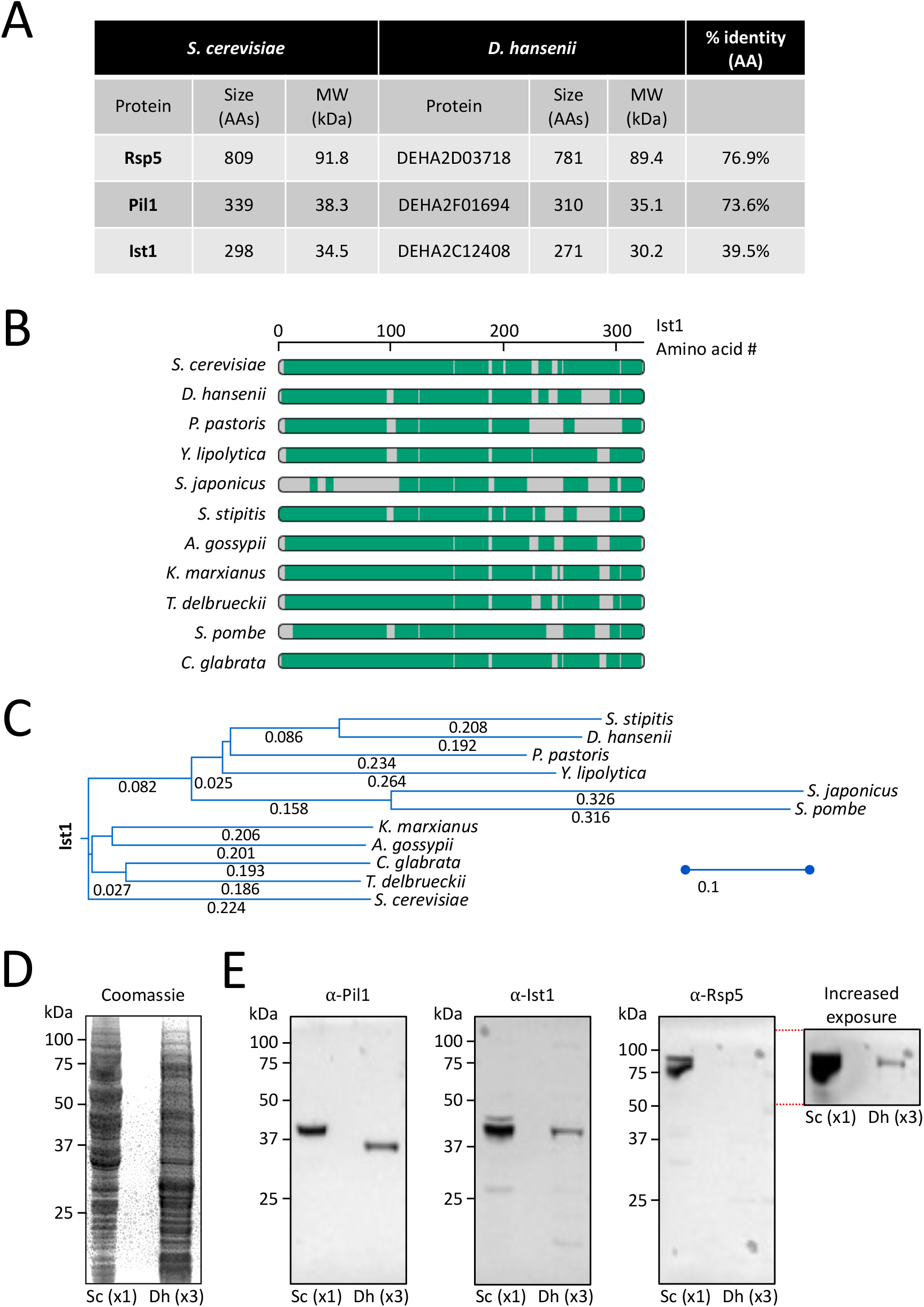
Proteomic comparisons between *S. cerevisiae* and *D. hansenii*. **A)** Table showing size of *S. cerevisiae* proteins Rsp5, Pil1 and Ist1 and their closest *D. hansenii* homologues identified by BlastP Search. **B)** Alignment of Ist1 protein sequence from the indicated yeast species. **C)** Phylogenetic tree of Ist1 from indicated protein sequences. **D)** Lysates were generated from equivalent volumes of yeast cultures and loaded 1x for *S. cerevisiae* and 3x *D. hansenii* cultures for SDS-PAGE followed by Coomassie staining (left) and immunoblot analyses with antibodies raised against Pil1, Ist1 and Rsp5 (right).

### Ultrastructural definition of Debaryomyces hansenii

Having compared *D. hansenii* to its better studied budding yeast counterpart *S. cerevisiae*, we next sought to optimise established tools from *cerevisiae* work to reveal details about the cellular architecture and dynamic processes in *D. hansenii*. The first structure we targeted for localisation was the nucleus. Nuclear stains have been used for many decades and are useful for demarcating individual cells, providing information on sub-nuclear concentrations of DNA, and for simple analysis of nuclear morphology and size. For this, we used Hoechst 33342, a blue fluorescent dye that binds to AT-rich DNA sequences (Latt and Wohlleb, 1975). Labelling of *D. hansenii* nuclei was very inefficient in SC media, therefore, we briefly incubated the cells in Tris buffer at pH 7.2 to increase nuclei labelling efficiency. Using this optimised protocol, the majority of cellular nuclei were defined (**Figure 4A**). Beyond this, focus on individual cells allowed clear indication of nuclear localisation that can be distinguished with vacuoles observable from DIC imaging (**Figure 4B**). These micrographs also demonstrate a small level of mitochondrial DNA separate from the nucleus. We next wanted to label mitochondria, which is highly important for metabolism of budding yeasts (Malina et al., 2018). For this, the carbocyanine-based dye, MitoTracker Green, label living *D. hansenii* cells in SC media (Chazotte, 2011). MitoTracker Green permeates the surface membrane and accumulates in active mitochondria (**Figure 4C**). As with *S. cerevisiae*, many cells exhibited a characteristic ribbon morphology with structures dispersed across mother and daughter cells during cell division (McFaline-Figueroa et al., 2011; Westermann and Neupert, 2000). Additionally, some other cells exhibited a more punctate localisation of mitochondria.

**Figure 4:**
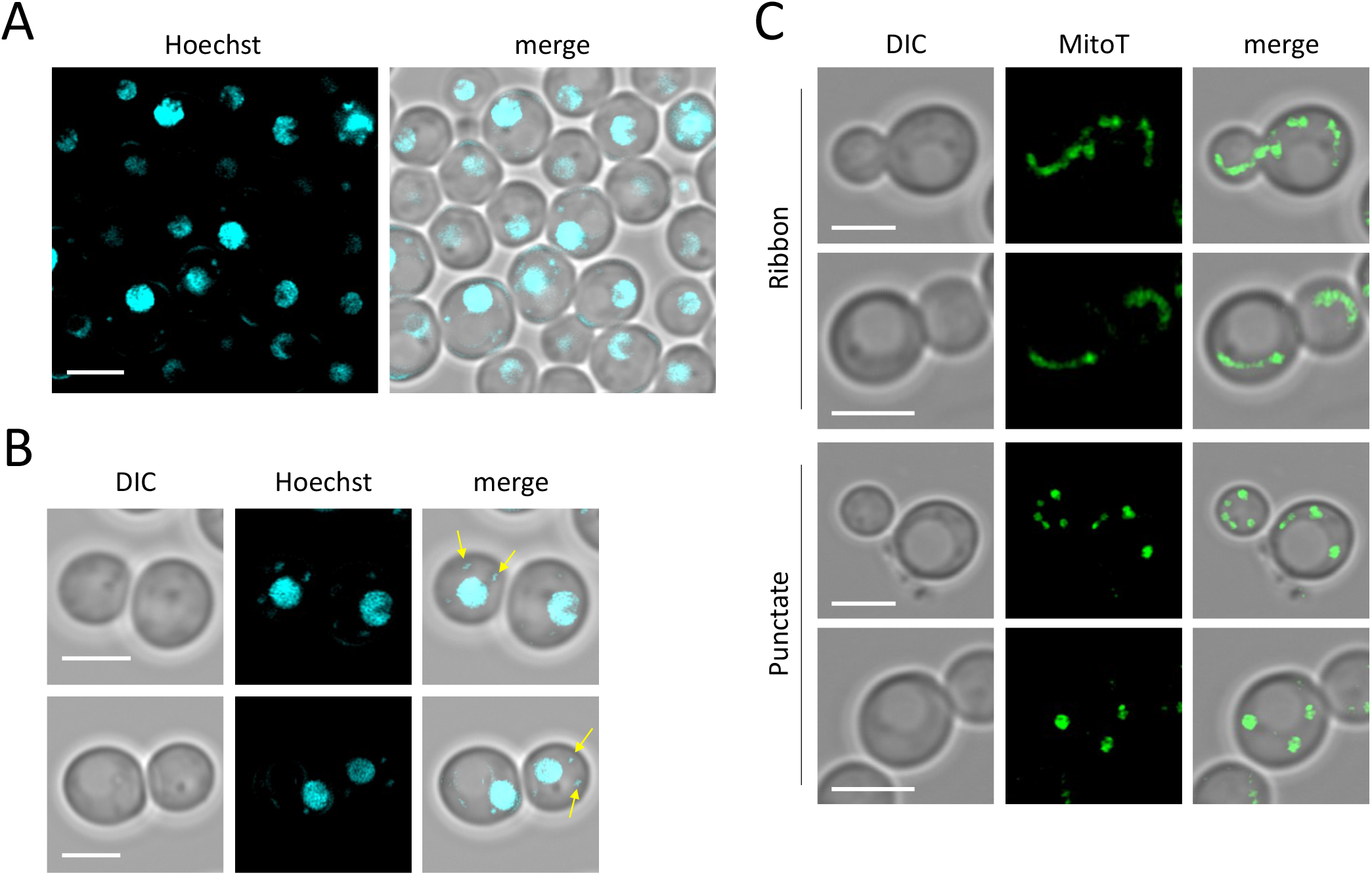
Imaging nuclei and mitochondria of *D. hansenii*. **A)** Log-phase cultures of *D. hansenii* were concentrated ∼10 fold, washed with Tris buffer at pH 7.2 and labelled with Hoechst 33342 for 30 minutes at room temperature. Cells were washed twice in buffer prior to confocal imaging using an Airyscan2 detector. **B)** Zoomed in images in same conditions as **(A)** to better highlight non-nuclear Hoechst staining (yellow arrows). **C)** Log-phase cultures of *D. hansenii* were stained with MitoTracker Green (MitoT) for 45 minutes in SC liquid media at 30°C prior to Airyscan2 confocal imaging. Scale bar, 3μm.

We next set out to define the cell wall of *D. hansenii* using Calcofluor-white (CFW), a blue fluorescent compound that binds cellulose and chitin of fungal cell walls and is enriched in chitin ‘bud-scar’ regions following separation of the budding daughter cell (Cabib and Bowers, 1971; Hayashibe and Katohda, 1973). As expected, CFW provides a bright signal at the cell wall, and allows bud scars to be observed (**Figure 5A**). Older cells exhibit more bud scars (Bartholomew and Mittwer, 1953), which was easier to assess by taking 45x 150nm z-slices, with individual scars only apparent at certain regions (**Figure 5B**) or from construction of 3D volumes (**Movie 1**). We also performed structured illumination microscopy (SIM) to obtain 3D images of CFW stained *D. hansenii* cells, to better highlight the discrete bud scars (**Figure 5C**). Finally, we tested the capacity of the blue dye CMAC (7-amino-4-chloromethylcoumarin) to label the vacuolar lumen of *D. hansenii* cells. Labelling of vacuoles was very efficient in minimal media, even following a brief incubation with CMAC (**Figure 5D**). The size and number of vacuoles per cell was easy to ascertain from CMAC staining, with fission-fusion vacuolar projections indicative of vacuolar dynamics observable (**Figure 5E**).

**Figure 5:**
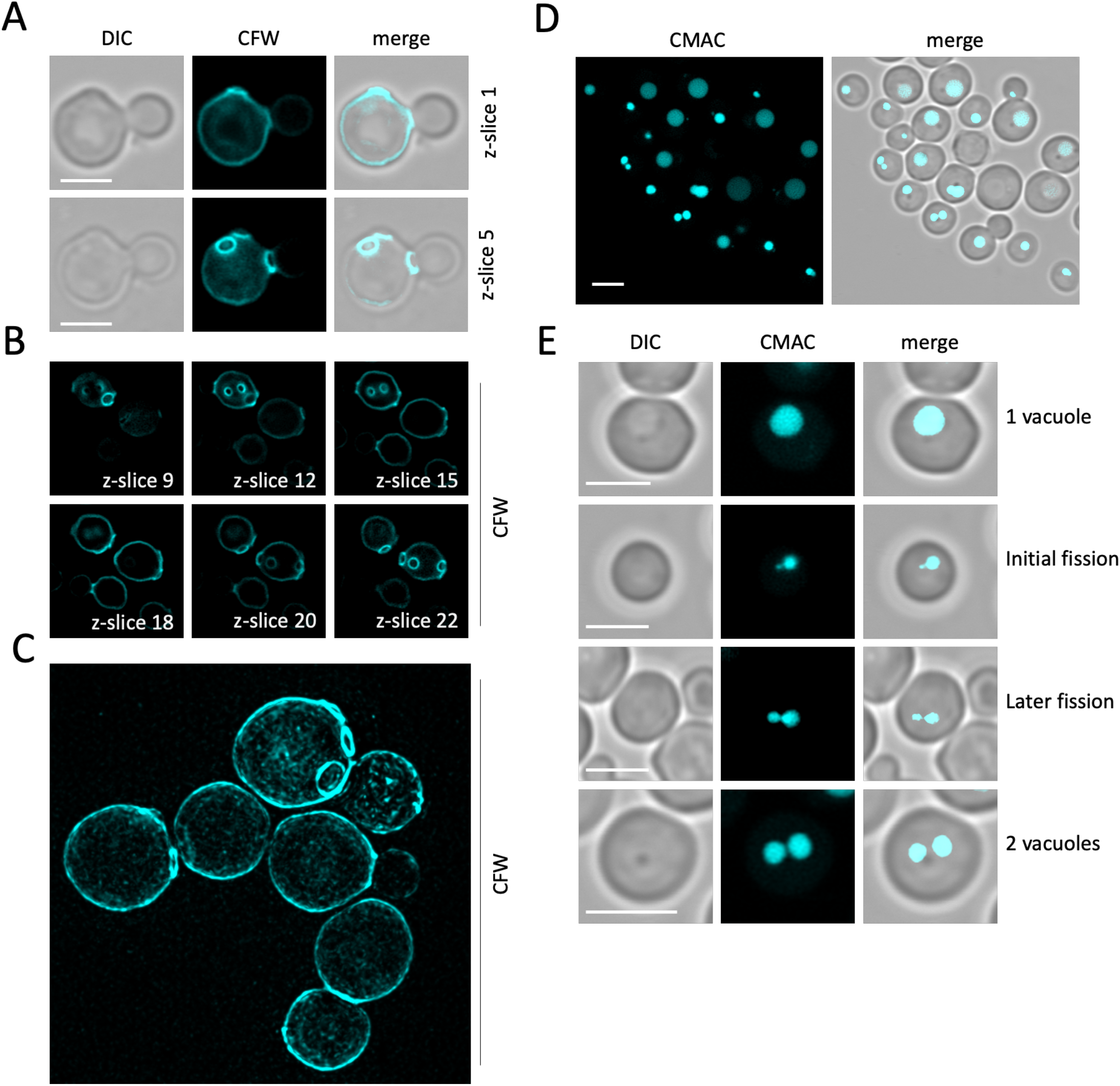
Imaging the *D. hansenii* cell wall and vacuolar lumen. **A - B)** Calcuflour white stain was added to cells in SC media for 5 minutes at room temperature prior to washing in SC media and preparation for imaging using Airyscan2. Individual z-slices are indicated to highlight different bud-scars across the volume of cells. **C)** Cells were labelled as above prior to lattice structured illumination microscopy followed by SIM^2^ processing, with an average intensity projection from all 42x z-slices shown. **D - E)** Cells were stained with CMAC dye for 15 minutes at room temperature followed by 3x washed in SC media and imaging using Airyscan2. Scale bar, 3μm.

### Studying organelle dynamics in Debaryomyces hansenii

To gain dynamic information about trafficking in the endolysosomal system, we used the styryl dye FM4-64, which is useful for visualising membranes in yeast, as it binds phospholipids of the plasma membrane and robustly increases fluorescence (Vida and Emr, 1995). We have recently employed an FM4-64 pulse chase technique combined with Airyscan2 to survey the endosomal system of *S. cerevisiae* (Andreev et al., 2023). A similar strategy was used to initially label the plasma membrane of *D. hansenii* cells, then temporally label endocytic trafficking to the lysosome / vacuole (**Figure 6A**). At first, the dye is exclusively localised to the surface membrane. Only 2-minutes of the label free chase was sufficient to observe significant endosomal signal in addition to the PM. Further time points show FM4-64 traffics from the PM to exclusively in endosomes at 10-minutes chase, followed by successful trafficking to the vacuole at 20-minutes and increasing over the remaining pulse period. This serves as a useful kinetic marker of endocytosis, in addition to spatially labelling the endolysosomal system of *D. hansenii* cells. We recently identified an endosomal recycling route that bypasses the *trans*-Golgi network in *S. cerevisiae* cells (Laidlaw et al., 2022b; MacDonald and Piper, 2017), efficiency of which can be measured by assessing the efflux of endosomal FM4-64 (Wiederkehr et al., 2000). We loaded *D. hansenii* endosomes with FM4-64 and measured efflux by flow cytometry, showing 58.3 ± 3.8% internalised dye is secreted via endosomal recycling in during the first 10 minutes (**Figure 6B**). We attribute this to active recycling mechanisms as the pre-treatment of FM4-64 endosome loaded cells with sodium azide, to inhibit active transport processes, entirely inhibits dye efflux.

**Figure 6:**
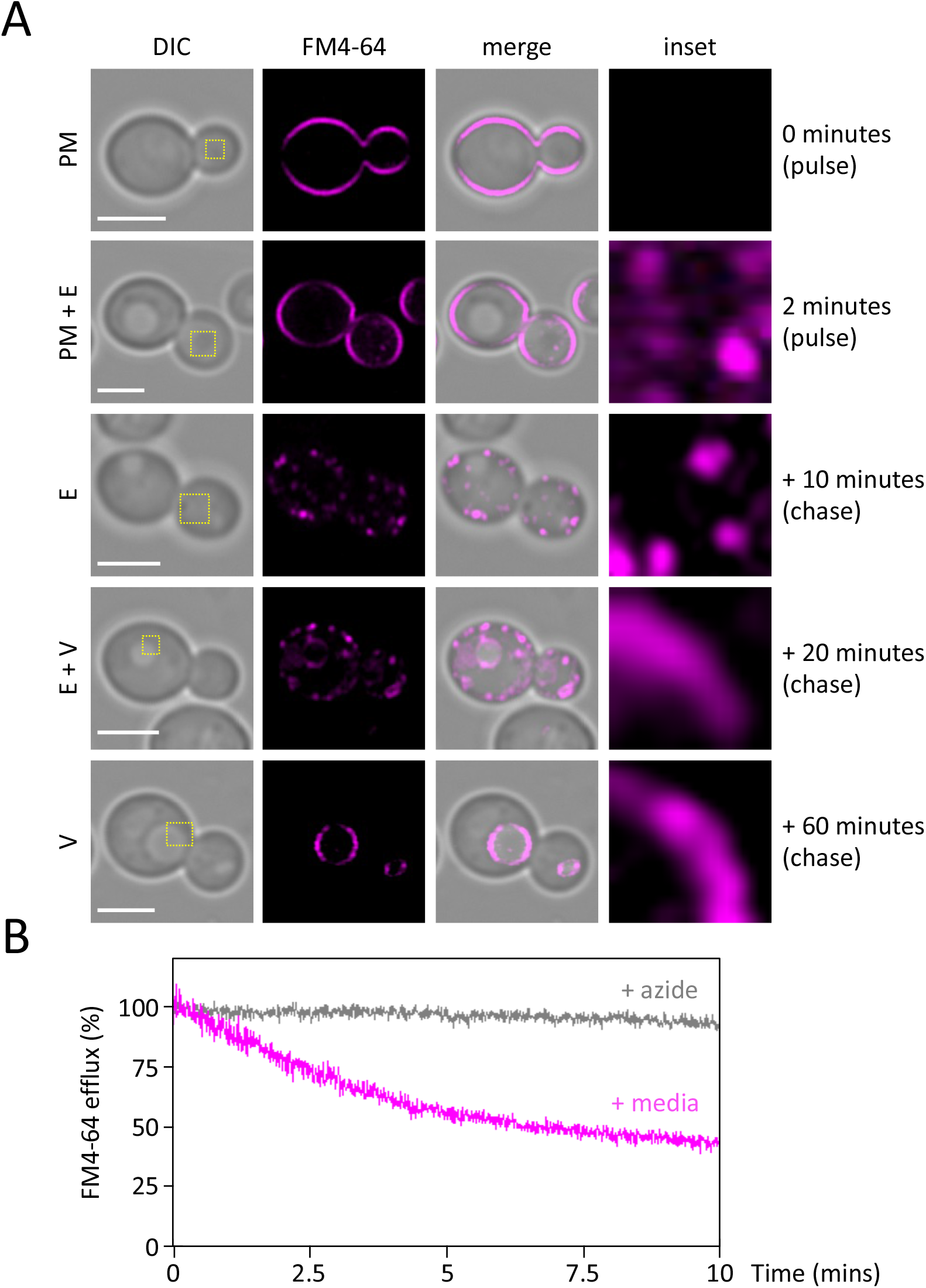
Visualising the endosomal degradation and recycling pathways in *D. hansenii*. **A)** FM4-64 labelling pulse chase was performed by addition of YPD media containing 8μM FM4-64 to harvest cells on ice (0 mins) followed by a 2-minute pulse incubation at room temperature (2 mins). Cells were then washed 3x 1 minute in ice cold SC media followed by incubation at 30ºC in SC media for +10-, +20- and +60-minute chase periods prior to harvesting. Localisations are labelled as plasma membrane (PM), endosome (E), vacuole (V). All time points were imaged by confocal Airyscan2 microscopy. **B)** Endosomes of *D. hansenii* were labelled in YPD media containing 40μM FM4-64 for 8 minutes followed 3x 3-minute washes in ice cold SC media. Fluorescence was then measured for 10-minutes following incubation in room temperature media (magenta) or azide kill buffer (grey). Scale bar, 3μm.

As holotomography was so efficient at rapid imaging entire cells, we next tested if intracellular refractive index (RI) contrast correlated with the vacuole. We expressed a GFP tagged nuclear marker in *S. cerevisiae* (GFP-Taf3) and imaged fluorescence simultaneously with holotomography measurements. Highlighting regions of particular RI contrast allowed the vacuole to be clearly distinguished from the GFP tagged nucleus (**Figure 7A**). Additionally, membranous elements of the plasma membrane and vacuole could be segmented separately. We then applied similar technique to image *D. hansenii* cells and this time, without a GFP integrated marker, we labelled the vacuoles with CMAC. Again, based only on refractive indices, the vacuoles of cells could be clearly observed (**Figure 7B**). This highlights the power of using holotomography as a label free technique, that can analyse specific organelles like the vacuole, for rapid imaging in live cells, without any prior genetic manipulations.

**Figure 7:**
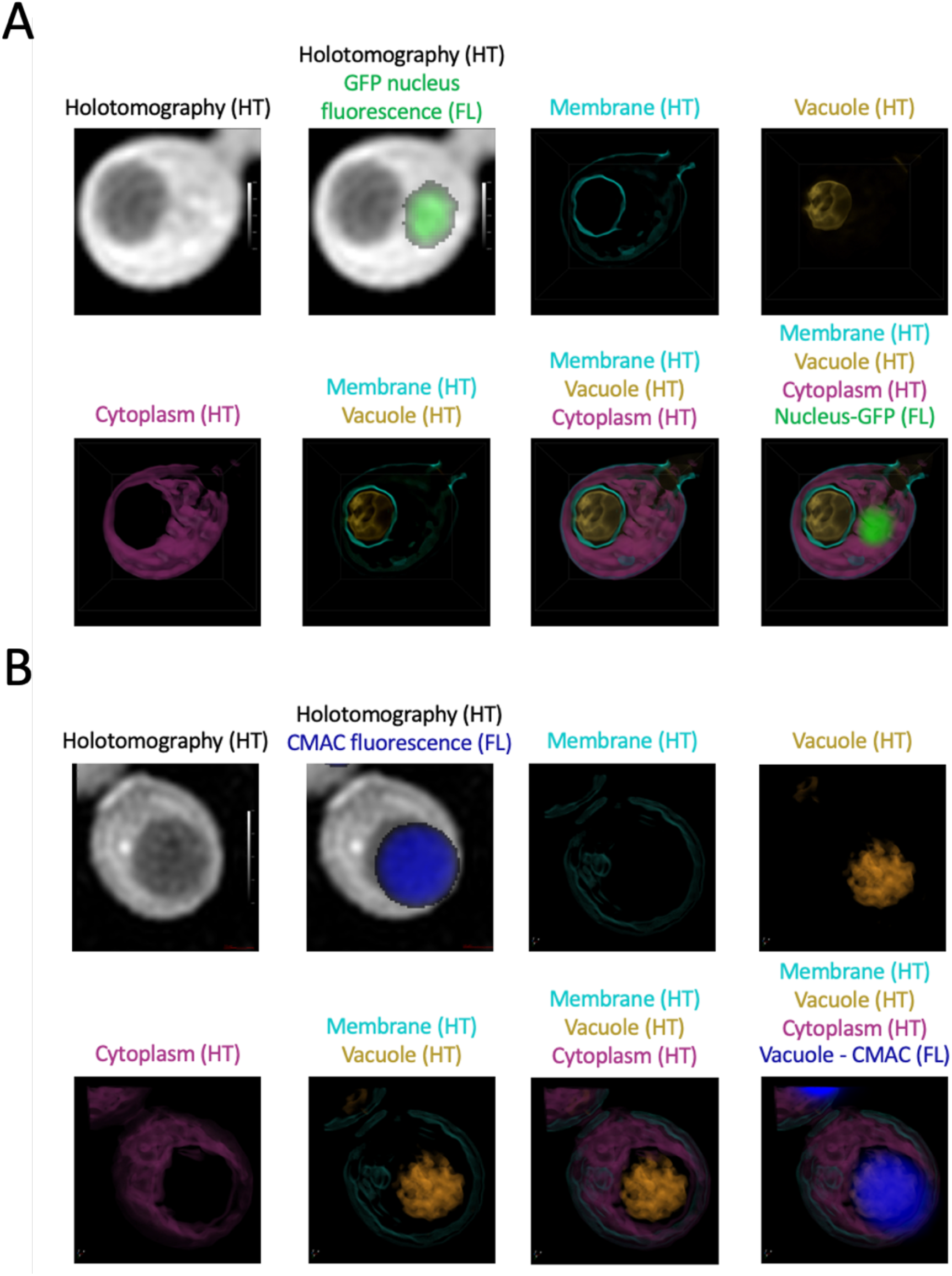
Optimising holotomography for label free visualisation of *D. hansenii* vacuoles. **A)** *S. cerevisiae* cells expressing GFP-Taf3 were cultured to log phase in SC media followed by preparation for holotomography (HT). GFP fluorescence (FL) was recorded and overlaid with raw HT. Additionally, indicated structures (membrane, vacuole and cytoplasm) were identified at specific refractive indices and visualised in blue, gold and purple, respectively. **B)** *D. hansenii* cells were cultured in minimal media prior to vacuolar labelling with CMAC and preparation for HT combined with CMAC fluorescence. As in **(A)**, refractive indices were adjusted to label individual aspects of cellular architecture.

Finally, organelles are inherited in budding yeast, with the sequential series of vacuole, mitochondria and nucleus being in part directed from the mother cell to the budding daughter cell in *S. cerevisiae* (Li et al., 2021). We therefore used the above-described methods to capture these inheritance events in *D. hansenii*. Using pulse-chased FM4-64, Hoechst, and MitoTracker, these dyes could be used to capture organelle inheritance events of the vacuole, nucleus and mitochondria, respectively (**Figure 8A - 8C**). Indicating that these biological events are also accessible to investigation in *D. hansenii*.

**Figure 8:**
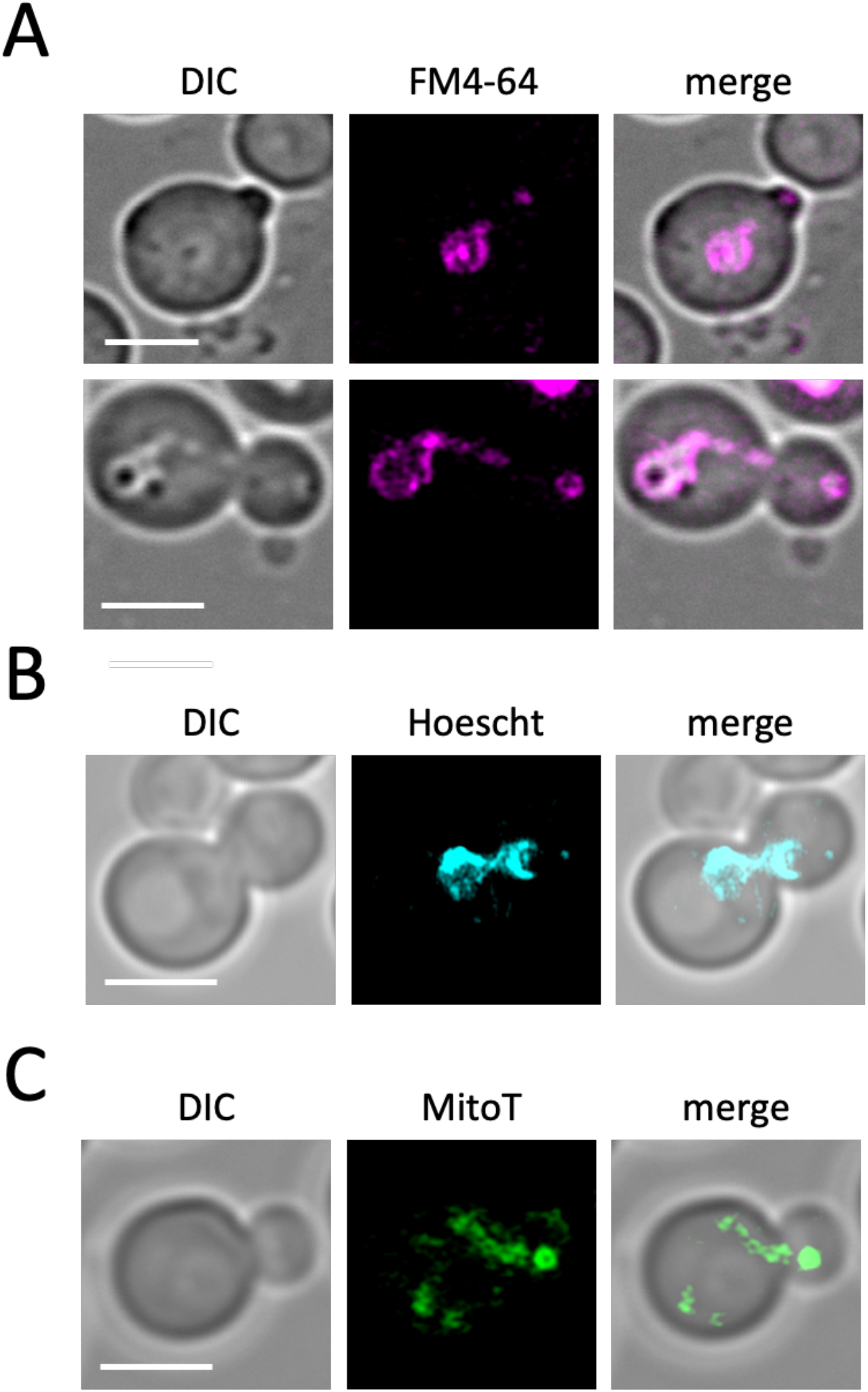
Using microscopy to study organelle inheritance in *D. hansenii*. **A)** Vacuoles of *D. hansenii* were labelled 8μM FM4-64 for 2 minutes followed by a 60 minute label free chase period in SC media. Extensions of labelled vacuoles were capatured by Airyscan2 confocal imaging at early (upper) and later (lower) I nthe bidding process. **B)** *D. hansenii* nuclei were stained with Hoechst in Tris buffer at pH 7.2 then prepared for Airyscan2 imaging. **C)** Log phase cells were imaged with MitoTracker Green in SC media prior to Airyscan2 imaging of mitochondria.

## DISCUSSION

There is growing interest in harnessing the biotechnological potential of the budding yeast *Debaryomyces hansenii*. For example, its extreme tolerance to salt and growth in alkaline conditions allows for low-cost sterile production in non-specialised conditions (Prista et al., 1997). *D. hansenii* metabolism has been shown as a valuable source for an array of products, such as xylitol, trehalose, D-arabitol, and several vitamins (Gírio et al., 2000; Nobre and Costa, 1985; Voronovsky et al., 2002). Beyond this, as *D. hansenii* also has a very high capacity to synthesize and accumulate lipids (Merdinger and Devine, 1965), it is an attractive microbial cell factory model system for production of both natural and non-natural chemical products (Breuer and Harms, 2006). Finally, *D. hansenii* can be used in fermentation processes, due to its osmotolerance and production of desirable flavour additives (Flores et al., 2004). Despite all these applications, little is known about the architecture of *D. hansenii* cells and study of its cell biology and biochemistry has been limited.

In this study we have made progress in the characterisation of *D. hansenii* by allowing observation of different biological features: 1) using live cell imaging combined with different fluorescent dyes; 2) employing label free holotomography; 3) utilization of antibodies generated against *S. cerevisiae* proteins for immunodetection of *D. hansenii* orthologues. Indeed, this study was initiated by comparison of *D. hansenii* with *S. cerevisiae*. Although these yeasts are relatively distant in evolutionary terms, and clearly exhibit different sizes and physical features (**Figures 1 - 2**), we did identify orthologues of *S. cerevisiae* proteins that were validated in *D. hansenii* lysates by immunoblotting (**Figure 3**). For example, the ESCRT-like protein required for endosomal recycling Ist1 (Laidlaw et al., 2022b), a pathway we were able to observe in *D. Hansenii* using a lipid dye efflux assay (**Figure 6**). Similarly, we were able to immunoblot the E3 ubiquitin ligase Rsp5, a master regulator of endosomal and vacuolar trafficking in *S. cerevisiae* (MacDonald et al., 2020; Sardana and Emr, 2021), which can be coupled to observations of endosomal trafficking kinetics by pulse chase experiments and vacuolar morphology observations using CMAC, FM4-64 and holotomography (**Figures, 4, 6 and 7**). Finally, although we were not able to visualise eisosome compartments using fluorescent methods, we were at least able to detect levels of the core eisosome factor Pil1. As Pil1 is post-translationally regulated in response to nutrient stress (Laidlaw et al., 2021; Paine et al., 2023), and eisosomes are generally associated with stress conditions (Athanasopoulos et al., 2019; Babst, 2019), it would be interesting to know whether eisosomes are involved in *D. Hansenii* salt tolerance. These are just some tractable inferences that can be made despite being unable to genetically modify and visualise trafficking pathways and biological processes. Similar approaches would be possible for any cross-species analysis, assuming there was sufficient conservation to monitor protein levels. We note that using antibodies raised against orthologues can lead to misinterpretation if comparing, for example, absolute levels across different species. Indeed, even the proteome profiles from lysates of *D. hansenii* and *S. cerevisiae* differ significantly (**Figure 3D**). However, these results do give an initial foothold to study such proteins in *D. hansenii* in different physiological conditions, where comparisons can be interpreted.

We also labelled several intracellular structures clearly with fluorescent dyes. For example, MitoTracker Green efficiently labelled yeast mitochondria in media, allowing for live cell imaging and morphological analyses (**Figure 4**). This is important, because modulation of respiratory activity in *D. hansenii* allows for adaptation to high salt conditions (Garcia-Neto et al., 2017). We also speculate that the high number of cells exhibiting fragmented or punctate mitochondrial morphologies, as opposed to the ribbon structures, might indicated that even our optimised culturing conditions may be inferior and stress inducing. Vacuole labelling was very effective, using both CMAC to label the vacuolar lumen and an FM4-64 pulse-chase protocol to label the limiting membrane of the vacuole (**Figures 5 - 6**). In the past we have struggled to effectively label *S. cerevisiae* nuclei with DAPI staining, resorting to fixation (Paine et al., 2021). Therefore, we tried various optimisation strategies using the DNA stain Hoechst 33342. This was successful with cells at different stages in the cell cycle having clearly identifiable nuclei (**Figures 4**). The cell wall staining procedure CFW was highly successful. This suggests the integrity of the *D. hansenii* cell wall could be ascertained through CFW treatment, as has been done for other fungal species (Hill et al., 2006; Ram et al., 1994). Beyond this, the chitin deposit scars created following asymmetrical divisions of budding daughter cells was easily visualised with different 3D imaging approaches, including Airyscan2 and lattice SIM (**Figures 5, Movie 1**). Although replicative life span can be assessed through micro-manipulation of diving cells (Kennedy et al., 1994) simply labelling with CFW and categorising cells based on how many scars are observed is a simple and effective method (Sinclair, 2013) now accessible to *D. hansenii* experiments. Unlike our other organelle labelling strategies in *D. hansenii* that allowed for efficient labelling and downstream imaging in media, staining of nucelli was only efficient in Tris buffer at pH 7.2. Although imaging of nuclei at steady state can provide various useful information, it is limited for live cell imaging in media. Although we can observe inheritance of the nuclei, only live cell imaging of FM4-64 labelled vacuoles and MitoTracker Green labelled mitochondria can be tracked in real time. In *S. cerevisiae* nuclear inheritance has been tracked by live cell imaging following transformation or integration of fluorescently tagged nuclear proteins, such as Nup59 or Nrd1 (Lecinski et al., 2024; Li et al., 2021). Therefore, it may be that further genetic developments will be required to convincingly track nuclear inheritance in *D. hansenii*.

In addition to fluorescence microscopy, we show holotomography based refractive index measurement, for example between the cellular contents and the extracellular media, works equally well in rich media compared to minimal media (**Figure S2**). Therefore, holotomography provides a very powerful imaging technique to complement other microscopy approaches, by allowing imaging in commonly used experimental media that is never typically used for imaging. In addition, holotomography allows for assessment of cells based on their stage in the cell cycle. We were able to ascertain whether cells were budding based solely on the sphericity measurements from the analysis software. Holotomography provides an exciting alternative for studying yeast species in conditions that mimic their natural environment, with the additional advantage of imaging cells very rapidly and without the requirement of tagging with fluorescent proteins, which can also potentially perturb the experimental system (Fei et al., 2011; Hoebe et al., 2008).

In summary, we have created a pathway to study many cell biological and biochemical aspects of the yeast species *D. hansenii*. The approaches we have employed, such as re-purposing antibodies, using fluorescent dyes, and harnessing label free imaging is a very powerful combination. Only a tiny fraction of known bacteria species are amenable to transformation (Johnston et al., 2014), systematic assessment of photosynthetic algae has only recently been achieved using a model species and most non-model species are inaccessible to investigation (Fauser et al., 2022; Mosey et al., 2021), and patient derived primary human cells are inherently limited due to the nature of their source and growth requirements (Richter et al., 2021). Therefore, arguably the most important biological cells - for human health and survival - remain inaccessible to scientific investigation and methods to overcome these challenges have high potential.

## METHODS

### Reagents

Wild-type strain BY4742 (Brachmann et al., 1998) was used for all *Saccharomyces cerevisiae* experiments apart from the GFP-Ycp4 strain (**Figure 1C**) and GFP-Taf3 (**Figure 7A**) tagged strains that are BY4741 background (Weill et al., 2018). The CB1 isolate of *Debaryomyces hansenii* (Ramos et al., 2017) was used for all other experiments.

### Cell culture

Both species of yeast cells were grown in either YPD (1% yeast extract, 2% peptone, 2% dextrose) or synthetic complete (SC) minimal media (2% glucose, 0.675% yeast nitrogen base without amino acids, plus appropriate amino acid dropouts for plasmid selection) (Formedium, Norfolk, UK). *S. cerevisiae* cells were routinely cultured at 30°C and *D. hansenii* cells were cultured at 25°C with shaking. Typically, a 2-fold serial dilution of was prepared and grown overnight so that mid-log phase cells could be harvested and used for experimentation the following morning.

### Microscopy

For microscopy, *S. cerevisiae* and *D. hansenii* cells from log-phase culture were labelled prior to concentration (∼10 fold) in SC media and then visualised at room temperature. For confocal microscopy, a laser scanning Zeiss LSM 980 equipped with an Airyscan 2 detector and a 63x / 1.4 NA objective lens was used. For Structured illumination microscopy (SIM), a Zeiss Elyra 7 system was used with a Plan-Apochromat 40x / 1.4 NA oil objective. Imaging in the blue channel for Hoechst 33342, Calcofluor-white, and CMAC was excited with a 405 nm laser (emission 422 - 497 nm), imaging in the green channel for GFP and MitoTracker-Green was excited with a 488 nm laser (emission 495-550 nm), and imaging in the red channel for FM4-64 was excited with a 561 nm laser (emission 570-620 nm). Airyscan2 and SIM^2^ processing was performed using Zen Blue (v3.4.1) and images were modified for brightness and pseudo-colouring using ImageJ software (NIH).

### Fluorescent labelling for microscopy

All *D. hansenii* cells were grown overnight to mid-log phase at 25°C prior to different labelling procedures. Where possible, labelling was performed in SC media, with the exception of Hoechst 33342 that only efficiently labelled yeast nuclei when labelling was performed in Tris buffer at pH 7.2. For nuclear staining, cells were concentrated ∼10 fold and washed once in Tris buffer at pH 7.2. The concentrated yeast sample was resuspended in 100μl buffer containing 2μl Hoechst 33342 from a 1mg/ml frozen aliquot. For mitochondrial staining, 200 μl of culture was incubated with 4μl MitoTracker Green stock at 100 μM concentration. For CMAC vacuolar staining, 200 μl of culture was incubated with 2μl CMAC from a 10 mM stock. For cell wall staining, 200 μl of culture was incubated with 2μl CFW, washed twice then imaged. FM4-64 pulse chase microscopy experiments were pulsed with 8μM dye for 2 minutes followed by label free media chases at indicated time points.

### Holotomography

Yeast cells were grown to log phase before imaging by holotomography at room temperature. Most experiments were acquired by 3D quantitative phase imaging (QPI) using an HT-2H instrument (Tomocube Inc), employing Mach–Zehnder interferometry and a digital micromirror device. 2D holographic images were acquired from 48 azimuthally symmetric directions with a polar angle (64.5°) controlled by a digital micromirror device (DMD) and used to construct a 3D tomogram based on refractive index. The diffracted beams from the sample were collected using a high numerical aperture objective lens (NA = 1.2, UPLSAP 60XW, Olympus) and recorded using a complementary sCMOS image sensor (Blackfly S BFS-U3-28S5M, FLIR Systems Inc.). For holotomography coupled to fluorescence imaging, of GFP-Taf3 *S. cerevisiae* and CMAC labelled *D. hansenii* cells, correlative 3D fluorescence images were acquired using the HT-2T instrument. Blue fluorescence data was acquired with an excitation filter of 392nm ± 12nm (381∼404 nm) and an emission filter of 432nm ± 18nm (414∼450 nm), while the green fluorescence was done with an excitation filter of 474nm ± 14nm (461∼487 nm) and an emission filter of 523nm ± 23nm (500∼546 nm). Semrock filters (IDEX Health & Science LLC) were used. The fluorescence z-stacks were acquired and processed with deconvolution software (AutoQuant by Media Cybernetics). The data was imaged and visualized using a commercial software (TomoStudio v3.3.9, Tomocube Inc., Korea).

### Flow Cytometry efflux assay

Mid-log phase *D. hansenii* cells were concentrated 10x in 200 μL YPD containing 40 μM FM4-64 dye (N-(3-Triethylammoniumpropyl)-4-(6-(4-(Diethylamino) Phenyl) Hexatrienyl) Pyridinium Dibromide) dye (Invitrogen™) to label endosomes for 8 minutes. Cells were then harvested in a pre-chilled centrifuge and supernatant fluid containi9ng FM4-64 discarded. Cells were then washed 3x in ice cold SC media. A concentrated sample of cold washed *D. hansenii* cells (∼20 μl) was then added to a flow cytometer tube and 3 ml of room temperature SC media at room temperature was added. Tube was immediately loaded on to a LSR Fortessa instrument (BD Biosciences) and fluorescence measurements from >1000 events per second were recorded for 10 minutes. FM4-64 fluorescence during cytometry was excited with a 561 nm laser, and data collected from an emission filter 710 - 750 nm. Data was visualised by converting fluorescence measurements to a percentage of the average fluorescence recorded in the first 10 seconds. Flow data were analysed using FCS Express (version 7.06.0015; De Novo Software).

### Immunoblotting

Log phase cultures of *D. hansenii* and *S. cerevisiae* were harvested, treated with 0.2 N NaOH for 5 minutes before resuspension in buffer (8 M urea, 10% glycerol, 50 mM Tris-HCl pH 6.8, 5% SDS, 0.1 % bromophenol blue and 10% 2-mercaptoethanol) to generate lysates. Lysates were resolved by SDS-PAGE then transferred to a nitrocellulose membrane using the iBlot2 transfer system (Invitrogen). Ponceau S and Coomassie staining was used to confirm successful transfer and equal loading, respectively. Rabbit polyclonal primary antibodies raised against Rsp5 (Stamenova et al., 2004), Pil1 (Walther et al., 2007), Ist1 (Tan et al., 2015) were individually incubated at 1/1000 concentration in 5% milk PBST (Phosphate-Buffered Saline, 0.1% Tween) overnight with rotation at 4°C. 6x 5 min washes in PBST were performed before incubation with 1/5000 IgG anti-rabbit secondary antibodies conjugated to HRP (Fisher Scientific Ltd) for 1 hour. Membranes were again washed 6x 5mins prior to visualisation. Membranes were visualised using a ChemiDoc Imager (Bio-Rad) in combination with enhanced chemiluminescence (ECL) Super Signal Pico Plus (Thermo).

### Data and statistical analyses

For cell area measurements, segmentation from brightfield micrographs was performed using the Cell Magic Wand Plugin for ImageJ. For holotomography analyses, cells were manually segmented and parameters isolated from the refractive index measurements using TomoStudio analysis Software v3.3.9 (Tomocube Inc). Unpaired Student’s *t*-tests were performed using GraphPad Prism v10.1.0 to compare the statistical significance between experimental conditions, with p-values included. An asterisk (*) is used to denote significance p < 0.05.

## ACKNOWLEDGMENTS

We would like to thank staff at the York Bioscience Technology Facility for technical assistance. This research was supported by a Sir Henry Dale Research Fellowship from the Wellcome Trust and the Royal Society 204636/Z/16/Z (CM).

## CONTRIBUTIONS

Conceptualization: CMD; Formal Analysis: MX, DG, CMD; Funding acquisition: CMD; Investigation: MX, DG, AM, SI, PM, GC, CMD; Project administration: JR, POT, PG, CMD; Supervision: POT, PG, CMD; Validation: MX, DG, AM, CMD; Visualization: MX, DG, AM, GC, CMD; Writing – original draft: MX, AM, JR, CMD; Writing

– review & editing: MX, DG, AM, JR, POT, PG, CMD.

## FIGURES & LEGENDS

**Movie 1: 3D volume of *D. hansenii* cell walls stained with Calcuflour white**

Log phase cells were stained with Calcuflour white in SC media for 5 minutes before washing and imaging in SC media. Airyscan2 confocal imaging was used to collect 45 individual z-stack slices, each 150nm. Zen blue was used to render a 3D projection across y-axis rotation and crop to indicated cells.

